# AI-assisted quantification of the clonogenic assay

**DOI:** 10.64898/2026.10.01.755964

**Authors:** Wilhelmina E. Radstake, Alexandra Kneringer, Janet M Denbeigh, Keerthy Rangan, Mahdi Bhalloo, Chris J. Beltran

## Abstract

The clonogenic assay is the foundational standard for evaluating cellular radiosensitivity, yet its manual quantification is limited by labor-intensive processing, high inter-observer variability, and a reductive binary scoring system. Here, we present and validate a deep learning-based segmentation pipeline for the automated, high-throughput quantification of mammalian cell survival across diverse radiation qualities (photons, protons, and carbon ions). By benchmarking against expert consensus using Bland-Altman analysis, we demonstrate that this AI framework consistently operates within natural human error margins. Crucially, transitioning to automated whole-well image-based analysis enables the extraction of continuous multidimensional data, including colony area, density, and cellularity. This phenotypic profiling successfully identified and quantified high-cellularity colonies formed by radioresistant BxPC-3 cells under particle irradiation. Coupled with rapid image acquisition platforms, this methodology significantly reduces processing bottlenecks while standardizing the derivation of relative biological effectiveness (RBE). By advancing beyond binary survival metrics to continuous morphological profiling, this framework provides a vital new analytical dimension for investigating structural mechanisms of radioresistance.

---

Clonogenic assays are a foundational methodology in radiobiology, used to assess the proliferative capacity of a single cell to form a colony following exposure to ionizing radiation. First described in the mid-20th century, this assay remains the gold standard for quantifying cellular radiosensitivity. The derivation of endpoints such as the surviving fraction (SF) and the application of radiation response models, specifically the linear-quadratic (LQ) model, provide critical parameters for both experimental and clinical radiobiology ^1,2^. Accurate and reproducible colony counting is therefore essential for generating reliable radiation dose–response curves and ensuring a meaningful interpretation of cellular response.

Despite their central role, the analysis of clonogenic assays remains a significant bottleneck in radiobiological research. Manual colony counting, which is still commonly employed, is time-consuming, labor-intensive, and subject to substantial intra- and inter-observer variability ^3,4^ .Variations in colony morphology, density, overlap, and imaging conditions further complicate consistent quantification. While several semi-automated tools and threshold-based image analysis pipelines (e.g., ImageJ macros, CellProfiler modules) have been developed, these approaches often require manual parameter tuning which involves a time-consuming process of trial-and-error and is vulnerable to suboptimal image segmentation, particularly when addressing the challenge of overlapping colonies ^5-7^.

Recent advances in artificial intelligence (AI), particularly deep learning-based image segmentation, offer new opportunities to overcome these limitations. Unlike traditional software that relies on rigid, user-defined thresholds, deep learning models can be trained to recognize underlying spatial patterns, making them far more adaptable to varying image conditions. Studies have explored the use of AI for colony detection, reporting improvements in automation and consistency ^8,9^. However, these investigations were primarily focused on bacterial colonies which exhibit relatively uniform circularity and distinct boundaries. In contrast, mammalian cell culture colonies present significant morphological heterogeneity and ill-defined borders which necessitates the development of a model that was specifically trained to handle these challenges.

In this study, we implement a deep learning-based segmentation framework as part of an image analysis pipeline aimed at improving the throughput, consistency, and objectivity of colony quantification for radiobiological applications. We focus on two morphologically distinct pancreatic cancer cell lines exposed to varying doses and radiation qualities. The results are evaluated against manual counting, and the segmented outputs are used to support downstream analyses relevant to preclinical radiobiology, including the determination of relative biological effectiveness (RBE).

## Methods

### Cell Lines and Culture Conditions

Two morphological distinct pancreatic cancer cell lines, PANC-1 and BxPc-3, were utilized in this study. All experimental specifics regarding cell culture, radiation exposure, dosimetry, and assay procedures are detailed in^10^.

### Manual Colony Quantification

Colonies containing 50 or more cells were defined as surviving and included in the count. Manual quantification was performed in a two-stage process: first, stained colonies visible to the naked eye were counted. The plates were then examined under a microscope to verify that all counted colonies met the 50-cell threshold and to correct for potential miscounts of borderline colonies. The X-ray datasets were counted by three independent observers, while the proton and carbon datasets were each evaluated by two observers.

### Automated Colony Quantification

Figure 1 outlines the methodological process of image acquisition, training of AI, and analysis of whole-well images.

**Figure 1.**
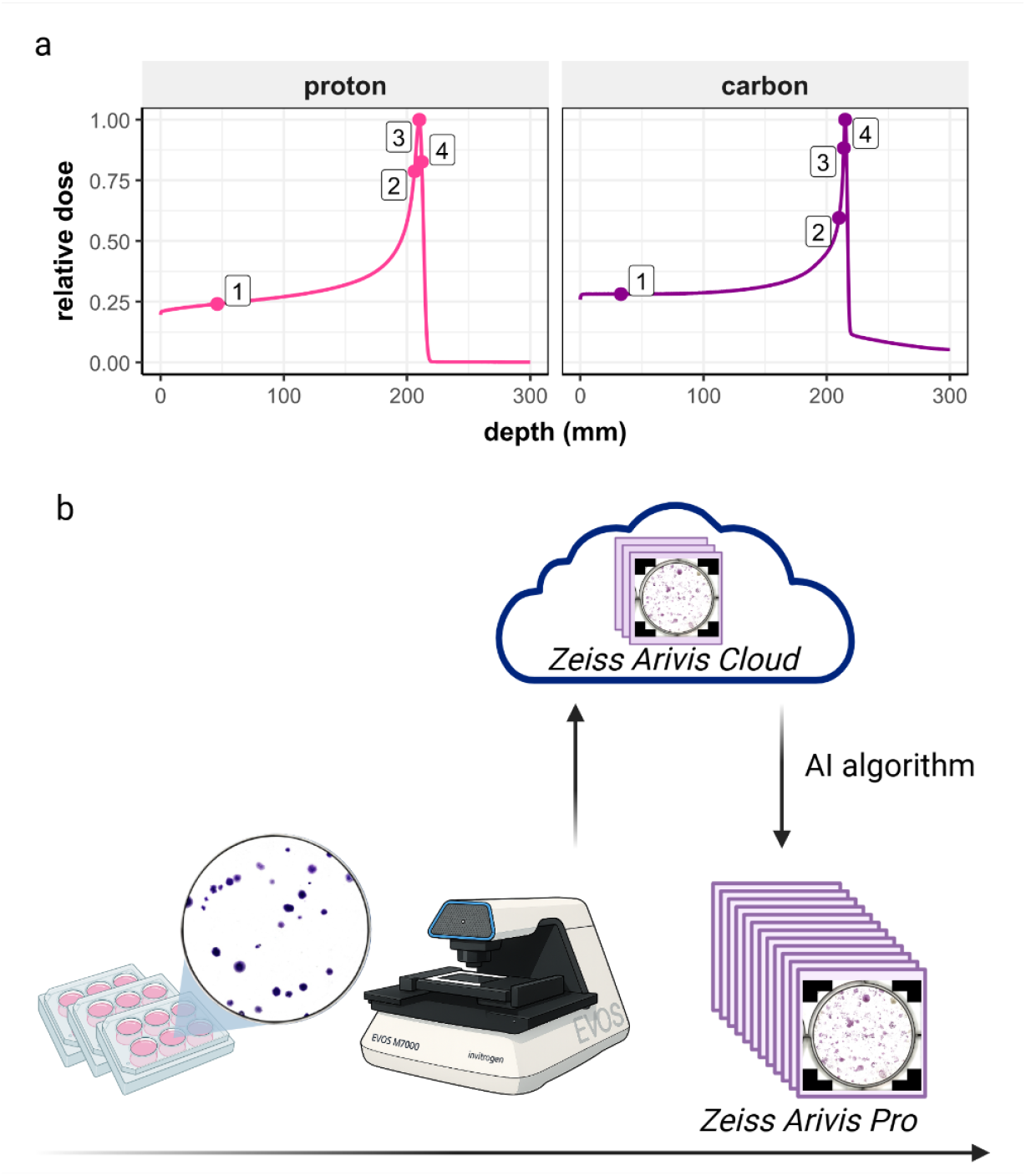
a) PANC-1 and BxPC-3 cells had previously been exposed to photon, and to proton and carbon ions at multiple positions along the Bragg curve. b) Cells were processed for analysis of clonogenic cell survival ^10^. Stitched whole well images were obtained using an EVOS M7000 equipped with a 2x objective and an 8-bit color camera. A subset of images (n = 17 for PANC-1, and n = 10 for BxPC-3 cells) were uploaded to Zeiss Arivis Cloud to train an AI segmentation model for each cell line separately. The final model was then downloaded and used for the segmentation step in an image processing pipeline, which was applied to all images in batch.

#### Image Acquisition

Whole-well images of crystal violet-stained colonies were acquired using an EVOS M7000 imaging system (Thermo Fisher Scientific, Waltham, MA) under brightfield illumination. Images were captured as 24-bit RGB files using a 2× objective lens, yielding a spatial resolution of 18.35 µm/pixel (1384 dpi). To capture the entire well area, individual fields of view were automatically tiled and stitched using the system’s default overlap settings. The stitched whole well tiff images were then saved for further processing.

#### AI Training

For each cell line, a subset of the acquired images representing a diverse range of colony morphologies was selected for training a deep learning-based segmentation model using the Zeiss Arivis Cloud platform. Whole-well images were uploaded and annotated using the built-in annotation tools to define the ground truth. Manual annotations were performed using a Wacom Cintiq Pro creative pen display, which allowed for precise digital outlining of colony boundaries.

The training process followed an iterative workflow, beginning with an initial round of approximately 50 independent colony annotations selected from across the entire dataset. Rather than labeling every object within a single image, specific colonies were chosen based on morphological variety to provide the model with a robust training set capable of generalizing across different experimental conditions. During this phase, colonies were defined by the classic radiobiological threshold of more than 50 cells, while non-cellular artifacts, media debris, and edge-of-well shadows were explicitly labeled as background to minimize false positives.

Once this baseline was established, the model was trained using the selected segmentation architecture. The resulting segmentation output was then qualitatively reviewed to identify discrepancies. Additional annotations were then added in subsequent iterations to refine the model’s performance. This iterative process of annotation, training, evaluation, and correction was repeated until the segmentation masks consistently matched the qualitative assessment of the raw images, ensuring sufficient segmentation quality for downstream analysis.

Initially, the study aimed to comparatively evaluate both semantic and instance segmentation architectures. However, iterative training of the instance segmentation model proved highly time-intensive and consistently yielded unsatisfactory results, failing to detect several colonies per well (false negatives) despite the addition of training images and annotations. Given that the primary objective of automating the workflow is to reduce the user intervention time, the instance segmentation approach was discontinued, and the study proceeded exclusively using semantic segmentation.

#### Image Processing Pipeline

The cell line specific semantic AI models trained in Arivis Cloud were imported into Arivis Pro (version 4.5.0). To account for the distinct morphological features of BxPC-3 and PANC-1 colonies, dedicated image analysis pipelines were developed independently for each cell line to quantify surviving colonies.

Within each respective pipeline, the corresponding semantic model was applied to generate an initial segmentation mask. Overlapping or closely positioned colonies were further separated using object-splitting algorithms. Parameters for split sensitivity were empirically optimized for each cell line to effectively separate true neighboring colonies while preventing artificial fragmentation. Object region growing was then employed to identify individual cells, and a parent-child relationship was established to associate individual cells (children) with their corresponding colonies (parents) (Figure 2). In accordance with standard clonogenic assay criteria, colonies containing fewer than 50 cells were filtered out and excluded from the final count. The finalized cell line specific pipelines were then applied in batch mode to process all corresponding whole-well images.

**Figure 2.**
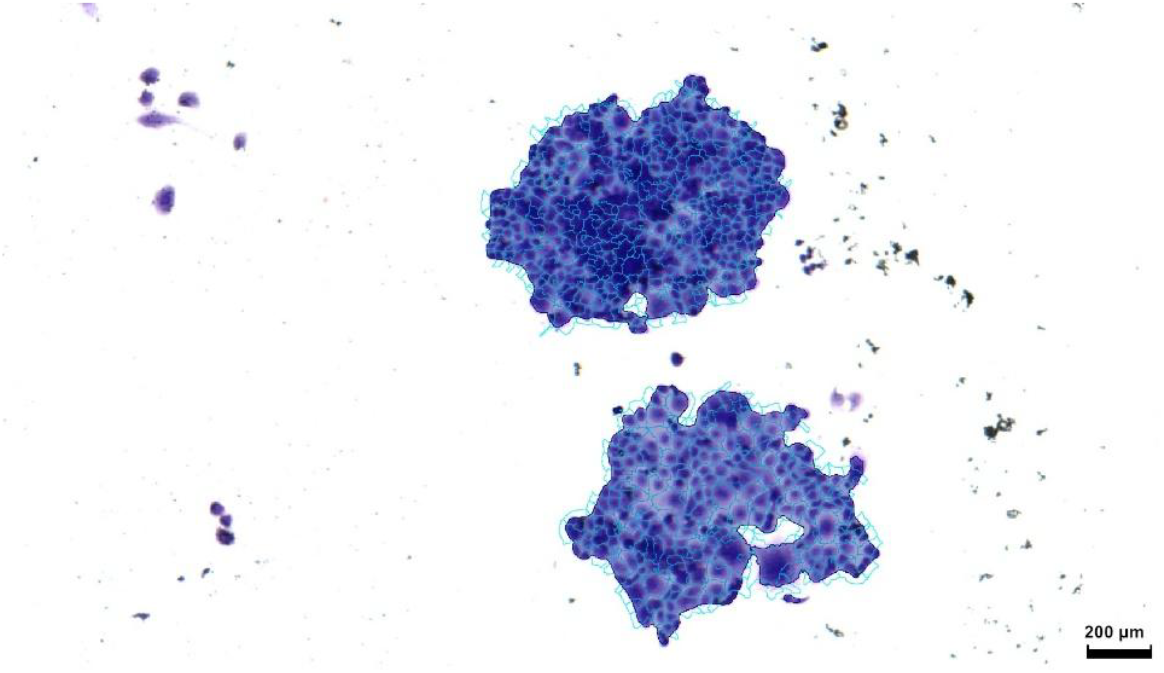
Example of segmented colonies and cells. Dark blue mask shows the colony as recognized by the AI algorithm. The cyan outlines the cells as detected by the seeded region growing step. Single cells and cell debris are classified as background by the AI algorithm and hence, not included towards the colony counts.

### Data processing

All analyses and visualizations were performed in R (Version 2025.09.1).

#### Modified Bland-Altman Analysis of Counting Agreement

To evaluate the performance of the automated counting models against inherent natural human variance, colony counts were compared between the AI pipelines and the consensus of trained human scientists. Standard Bland-Altman limits of agreement (LoA), which evaluate absolute counting differences, are susceptible to heteroscedasticity in clonogenic assays, as absolute variance typically scales with total colony density. Therefore, the percentage difference was employed as the comparative metric.

For each unique experimental condition, defined by cell line, radiation quality, physical position along the Bragg peak, dose, and replicate, the mean of the independent human observers was calculated as the reference consensus. Any conditions missing valid counts from at least two independent human observers were excluded from the variance analysis. The relative counting error for each individual human observer, as well as for the AI models, was then calculated as the percentage difference from this reference:

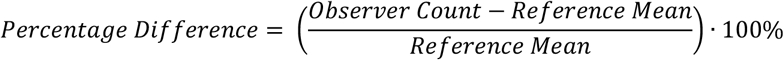

The percentage differences were plotted against the reference mean to visualize proportional error across the colony density spectrum. To quantitatively assess this agreement, the mean systematic bias and the 95% LoA were calculated for each cell line and observer type. The mean bias was defined as the average of the percentage differences, while the 95% LoA were calculated as the mean bias ± 1.96 x SD (standard deviation of the percentage differences). These human-derived error boundaries served as a benchmark for evaluating the AI models’ performances.

#### Survival Fraction and Relative Biological Effectiveness

Survival fractions were calculated normalizing each plating efficiency (PE) of irradiated samples to the mean PE of the corresponding unirradiated control group. The parameters α and β of the linear-quadratic model (LQM) were estimated by fitting the log-transformed experimental data to the equation:

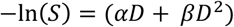

Fitting was performed using a non-linear least squared (nls) approach in R. Specifically, the “port” algorithm was employed to enforce non-negative constraints to avoid biological nonsensical upward curves, with starting values set at 0.5 Gy^-1^ for α and 0.01 Gy^-2^ for β. Model parameters were extracted for each observer group. Observers were categorized as either “AI Model” or “Human Observer” to facilitate direct comparison of the fitted parameters.

To represent experimental variance and model uncertainty, a multi-tiered statistical approach was implemented. For raw data points, the mean survival fraction and standard deviation were calculated for each experimental condition. To account for the uncertainty in the LQM fits, confidence intervals for the α and β coefficients were derived from the standard errors (SE) obtained during the non-linear least squares optimization. Upper and lower survival bounds were calculated by applying the 95% confidence interval (±1.96 x SE) to the parameters. The upper survival bound was calculated by using the minimum coefficients (α_min_, β_min_), while the lower survival bound utilized the maximum coefficients (α_max_, β_max_):

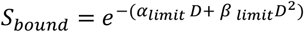

A human consensus band was constructed to define the aggregate uncertainty of the manual counting cohort. This band was defined by the absolute minimum and maximum survival values across the confidence envelopes of all human observers at each dose point. The resulting envelope represents the “gold standard” variance, incorporating both the inter-observer counting differences and the statistical uncertainty of the resulting model fits. The AI-derived survival curves and their associate confidence intervals were then evaluated against this consensus band to determine methodological agreement.

The estimated α and β for all observers and radiation qualities were then used to calculate RBE values for the surviving fraction of 10% using the equation (Parisi et al 2020):

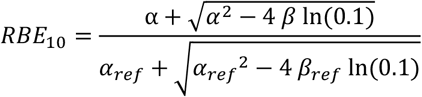

Where α, β, α_ref_, β_ref_ are the linear and quadratic terms of the LQM for the radiation under investigation and the reference photon (X-ray) exposure respectively.

The RBE_10_ for each human observer was calculated using their own specific photon reference data. Similarly, the AI-derived RBE_10_ was calculated using the reference generated by the AI model.

For comparative analysis, the human consensus range was established for each experimental condition, defined by the mean, minimum, and maximum RBE_10_ values across the human cohort. The AI models’ performances were evaluated by calculating the ratio of the AI-derived RBE to the human mean RBE, allowing for direct assessment of whether the automated pipeline fell within the natural range of human inter-observer variability.

## Results

### Quantitative Bland-Altman Agreement

To quantify counting performance of the AI algorithms, a percentage-based Bland-Altman analysis was conducted on the pooled reference data for each cell type (Figure 3, Table 1).

**Table 1.**
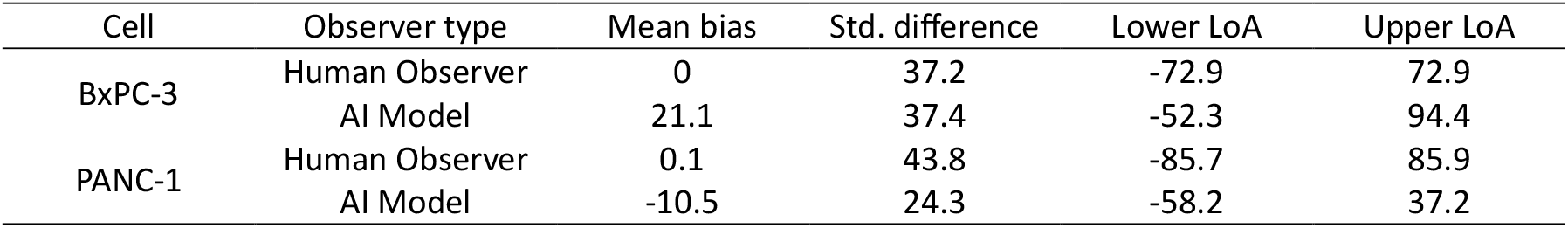
Bland-Altman analysis of colony counting agreement.

**Table 2.**
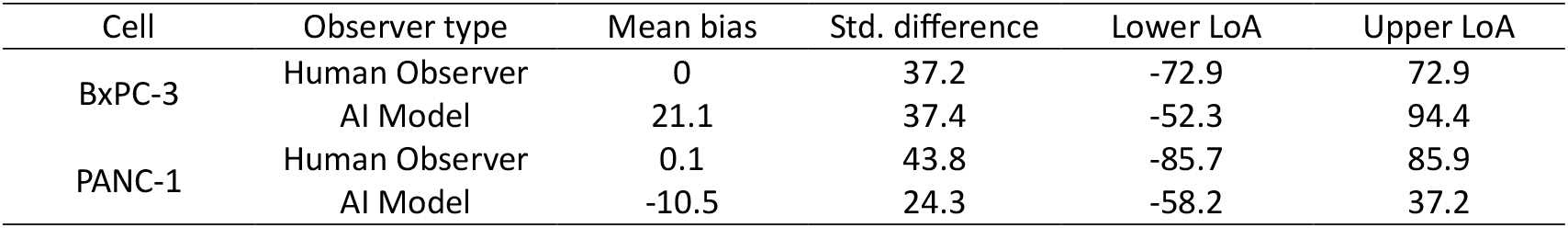
Bland-Altman analysis of colony counting agreement.

**Figure 3.**
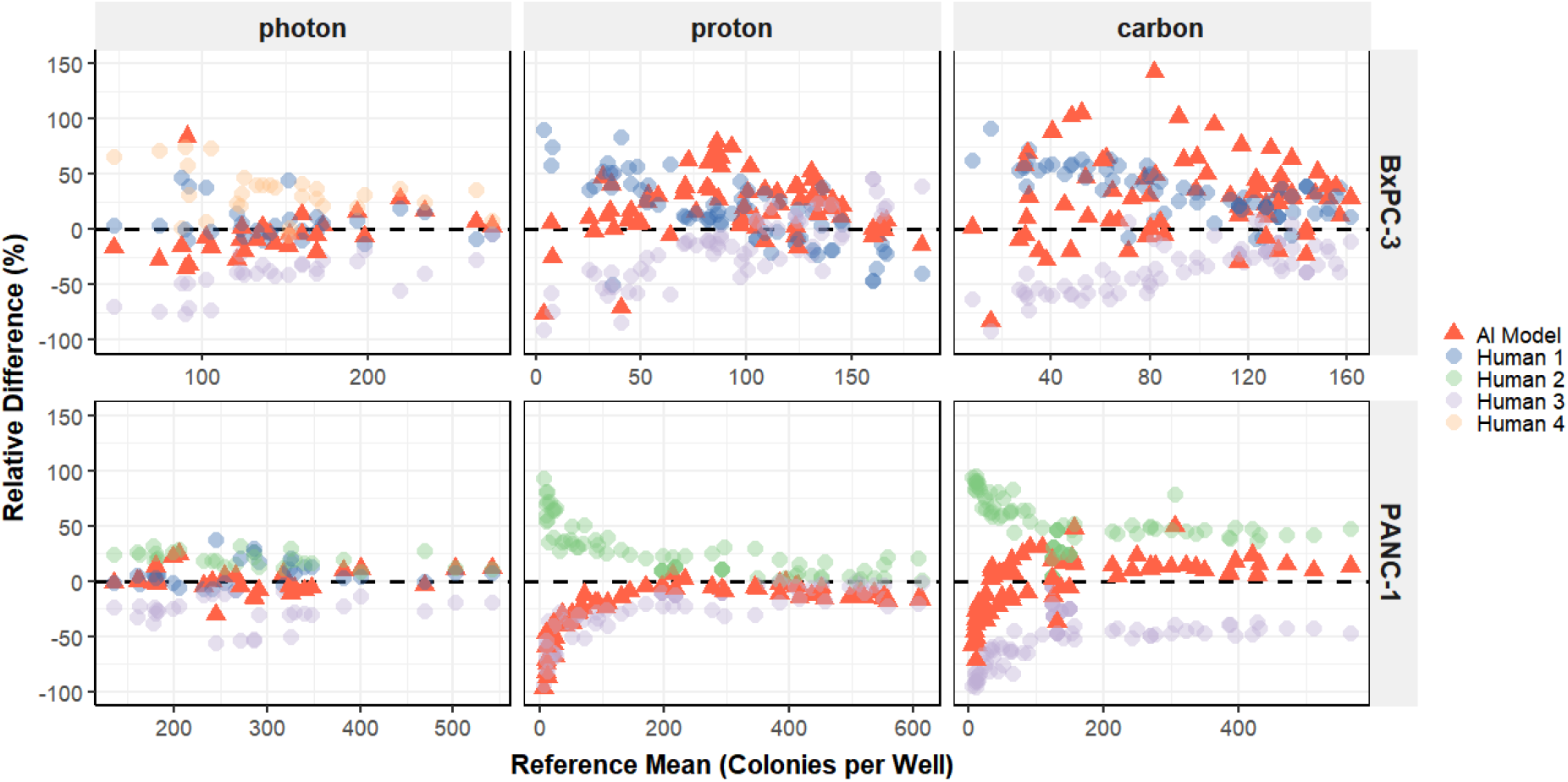
Agreement analysis of automated AI colony counting versus human consensus across radiation qualities. Modified Bland-Altman plots display the relative percentage difference in colony counts between individual observer and the human reference mean (x-axis) for BxPC-3 and PANC-1 cell lines. The horizontal dashed black line represents perfect agreement (0% difference). Individual human observers (circles) demonstrate natural inter-observer variability clustered around the zero line. The AI model’s predicted counts (red triangles) demonstrate high concordance with human observers in PANC-1 across all modalities and in BxPC-3 under photon irradiation. A bias towards higher counts for the AI model is observed under proton and carbon irradiation for BxPC-3 cells.

For the PANC-1 cell line, the AI model demonstrated good agreement with the human reference mean. While a slight systematic undercounting bias of 10.5% was observed for the AI model, the 95% limits of agreement (LoA) ranged from -58.2% to 37.2 %. This variance was remarkably narrower than the natural human baseline variance, which had a 95% LoA of -85.7% to 85.9%. This confirms that for the PANC-1 cell line, the AI model was consistently more precise and consistent than the human baseline, effectively reducing the inherent inter-observer variability typically found in manual colony counting.

For the BxPC-3 cell line, the AI model exhibited a systematic overcounting bias of 21.1%, shifting its 95% LoA to a range of -52.3% to 94.4% compared to the human reference 95% LoA of -72.9% to 72.9%. However, the standard deviation of the AI’s differences (37.4 %) was virtually identical to the human pooled standard deviation (37.2 %). This indicates that while the AI’s systematically produces higher total count, the precision and reproducibility of the model match human baseline variance.

To further investigate the overcounting bias for the BcPC-3 AI model, two representative experimental conditions were isolated: one demonstrating high agreement with the human reference mean (photon, 0Gy; +2.28% relative difference) and one with high over estimation (carbon, Bragg peak, 2 Gy; +142% relative difference).

Visual inspection of the whole-well images reveals a phenotypic difference between the two conditions. In un-irradiated control wells, BxPC-3 colonies present as distinct, relatively uniform clusters that are distributed across the well (Figure 4a). In contrast, high-LET carbon irradiation induces morphological heterogeneity. Whole-well images show irregular colony morphologies, characterized by large cell clusters mixed with smaller fragmented colonies (Fig 4b). The density distribution of number of cells per colonies between the two conditions further supports this observation (Fig 4c). In the control condition, most colonies contain about 100 – 300 cells. However, the carbon irradiated wells show a peak of colonies with fewer than 100 cells, in addition to a large tail extending out to 10,000 cells per colony.

**Figure 4.**
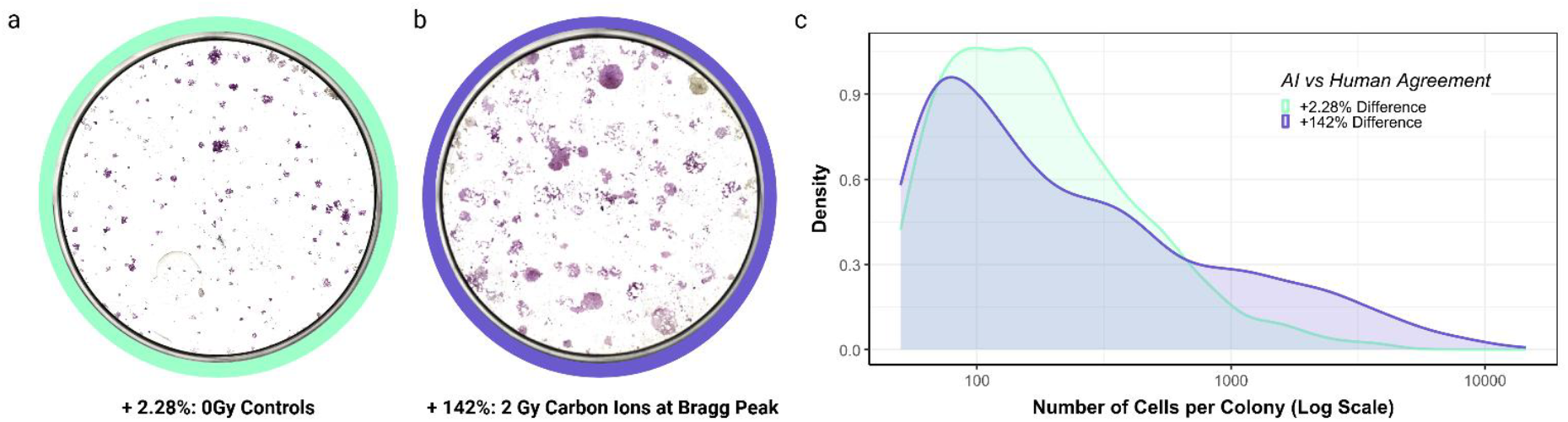
(a) Representative whole-well image of a BxPC-3 control condition, demonstrating uniform colony morphology and high agreement (+2.28% difference) between the AI model and human consensus. (b) Representative whole-well image of BxPC-3 colony formation after exposure to 2 Gy carbon ions at the Bragg peak, demonstrating irregular, clustered colonies mixed with smaller colonies, corresponding to an AI overcount (+142% difference) relative to human observers. (c) Logarithmic density distribution of cells per colony for the corresponding conditions. The 0 Gy control (green line) exhibits a peak between the 100 – 300 cells per colony. The 2 Gy carbon condition (purple line) exhibits a dual distribution, maintaining a peak near <100 cells per colony while developing a large tail extending to over 10,000 cells.

**Figure 5.**
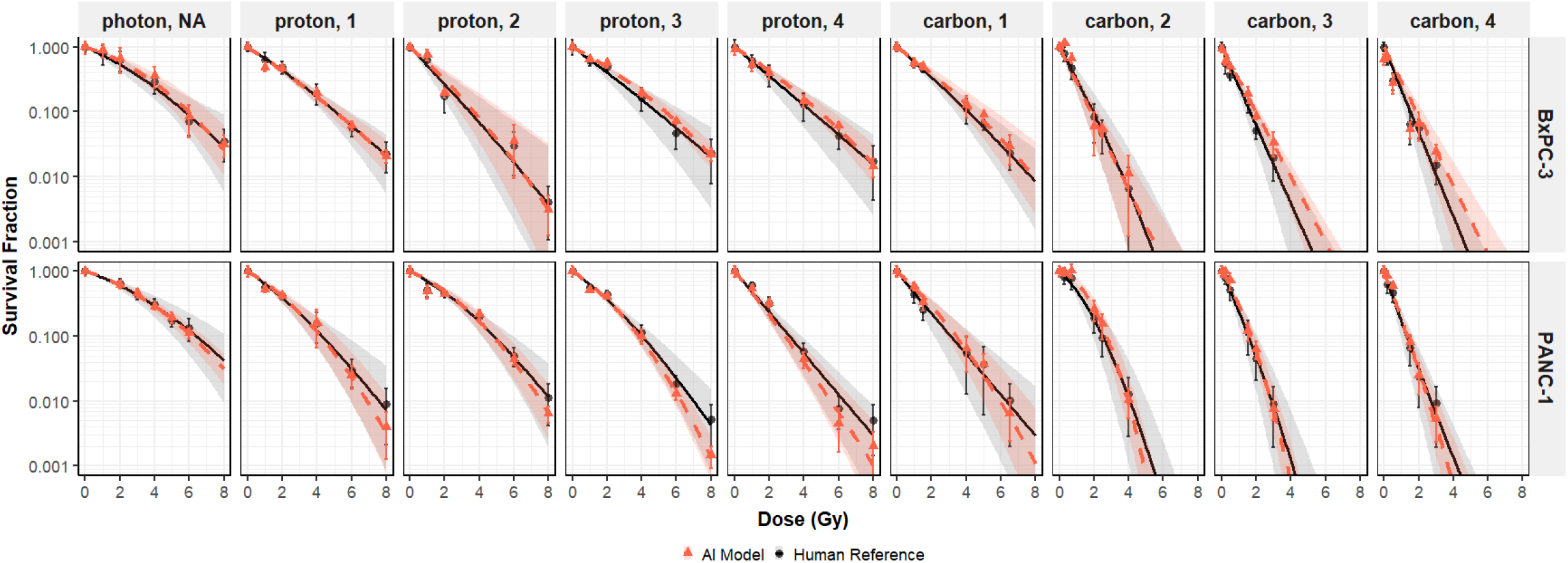
Comparative clonogenic survival of BxPC-3 and PANC-1 cells across radiation qualities. Clonogenic survival fractions are plotted against physical dose (Gy) for photon irradiation and proton and carbon ions at designated Bragg curve positions. Grey circles and orange triangles represent the mean survival fraction at each dose as determined by the human reference mean and the automated AI model, respectively. Error bars represent the standard deviation across independent biological iterations. The dashed orange line represents the fitted linear-quadratic survival curve derived from the AI model’s counts. The solid black line represents the corresponding fit derived from the human reference data. The broad grey shaded region illustrated the absolute bounds of natural human inter-observer variability, capturing the minimum and maximum survival fractions calculated by independent human scientists. The orange shaded region represents the statistical uncertainty bounds of the AI model’s predicted fit.

This suggests that the presence of large colonies may negatively influence the observer’s accuracy by causing humans to systematically overlook or subconsciously discount the smaller, paler colonies that still clear the 50-cell survival threshold. Since the AI-model performs a pixel-by-pixel evaluation, this threshold is uniformly applied to all identified colonies, resulting in higher overall counts.

#### Survival Fractions and RBE

Next, to evaluate how differences in raw colony counts between human observers and the AI pipelines influence further downstream analysis, we calculated the survival fractions for each experimental condition and fitted survival curves using a standard linear-quadratic model (Fig 4).

For the PANC-1 cell line, the survival curves generated by the AI model demonstrated close alignment with the human reference curves across all radiation conditions, and the AI-derived survival fractions consistently mapped within the bounds of human inter-observer uncertainty. This supports the model’s accuracy and reliability for determining survival of PANC-1 cells exposed to varying radiation qualities.

In the BxPC-3 cell line, for the photon and proton exposed cells, tight agreement between AI and human-derived curves was observed. However, the bias of the AI model to higher colony counts resulted in deviation of the fitted AI curves from the human reference curves, yielding systematically higher survival fractions which were especially visible at the higher-LET radiation conditions.

We next calculated the relative biological effectiveness (RBE) at 10% survival for all conditions and observers. The RBE values generated using the AI analysis pipelines demonstrated substantial agreement with the human observers (Fig 6a). Both observers accurately captured the expected rise in RBE as a function of penetration depth, reflecting the characteristic increase in linear energy transfer (LET) as the proton and carbon ion beams traverse deeper into the water phantom. To quantify the AI’s performances relative to the human disagreement, the data were normalized to the human reference mean (Fig 6b). This comparative evaluation showed that both AI models’ relative deviations consistently remained within or closely adjacent to the human interobserver variability. Notably, the most pronounced deviations occurred in the BxPC-3 cell line at the carbon ion positions 3 and 4 where RBE10 values were more than 10% lower than the human consensus, which is in concordance with the earlier observed increase in raw colony counts in those conditions.

**Figure 6.**
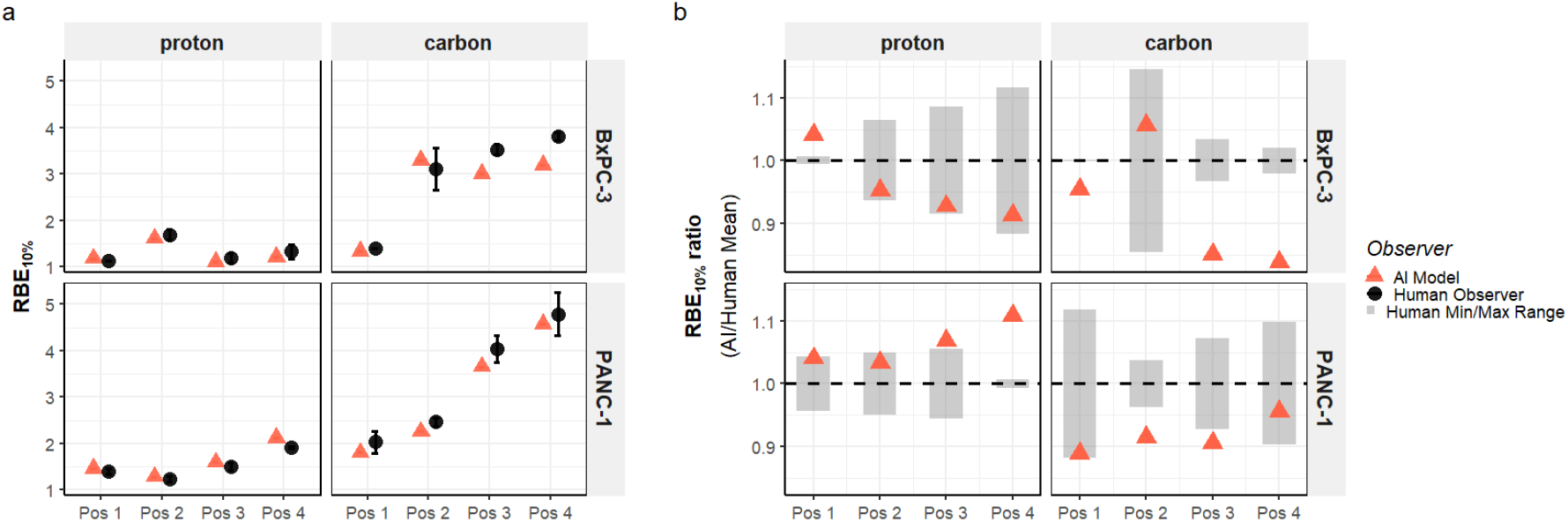
RBE10 values (a) and ratio’s (AI/Human) (b) are plotted for BxPC-3 (top row) and PANC-1 (bottom row) cell lines exposed to proton (left column) and carbon ion (right column) radiation as function of physical positions along the Bragg curve (Pos 1-4). Error bars represent the absolute minimum and maximum RBE10 values calculated independently by the two human observers, illustrating inter-observer variability. The dashed horizontal line at y = 1.0 represents perfect agreement. Grey shaded rectangular regions represent absolute biological uncertainty, defined by the minimum and maximum RBE10 ratio limits established by the individual human experts.

#### Colony cellularity

In addition to obtaining objective quantification of cell survival, automated analysis of clonogenic images enables extraction of quantitative features that were previously unmeasured. To explore this novel capability, we next examined the relationship between radiation quality and colony growth. We quantified the average cells per colony and by mapping this against the standard survival curves, we identified two distinct cell responses to particle therapy (Fig 7).

**Figure 7.**
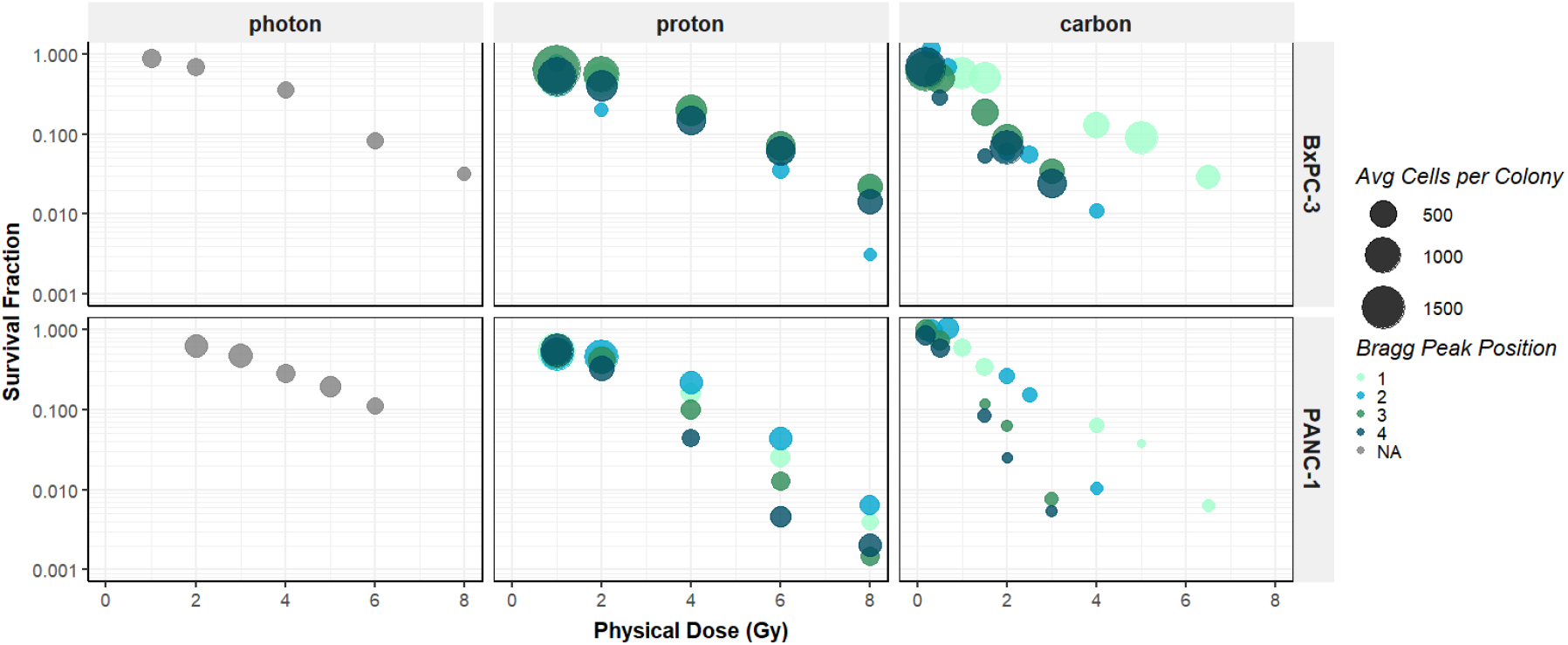
Clonogenic survival fractions are plotted against physical dose (Gy) for photon irradiation and proton and carbon ions at multiple Bragg curve positions for BxPC-3 (top row) and PANC-1 (bottom row) cell lines. The size of the circles represents the average cells per colony for the specified radiation quality and dose point.

In the PANC-1 cell line, we observed a characteristic proliferation-inhibition paradigm. As survival decreased with increasing dose, the average cells per colony remained stable or as observed after carbon exposure, decreased. Contrary, the BxPC-3 cells did not show this trend and an increase in average cells per colony after proton and carbon as compared to photon exposure was observed.

## Discussion

The primary objective of this study was to validate a deep learning-based image analysis pipeline for the automated quantification of the clonogenic assay. Automating this process is highly desirable as the manual counting of colonies remains widely applied but is notoriously labor- and time-intensive. In this study, human variability served as error margins for benchmarking our AI segmentation models. Our main findings demonstrate that AI’s performance consistently fell within these established boundaries. These results validate the application of automated segmentation for quantifying clonogenic cell survival across two morphologically distinct mammalian cell lines, effectively standardizing the derivation of downstream radiobiological parameters.

A key finding of this work is the observation of a distinct shift in colony morphology following exposure to proton and, particularly, carbon ions in the previously reported radio-resistant BxPC-3 cell line ^10^. A primary limitation of traditional manual scoring is its reliance on a binary threshold, which reduces complex cellular responses to a simplistic “survivor” or “non-survivor” classification. By transitioning the analysis to whole-well images, our framework extracts a variety of objective, quantifiable metrics such as colony area, density, and cellularity. Using this multi-dimensional data, we were able to detect and quantify the large, high-cellularity colonies uniquely formed by BxPC-3 cells under particle irradiation. Capturing these structural adaptations provides critical new insights that could serve as the foundation for future studies investigating the underlying mechanisms of radioresistance.

Notably, the Bland-Altman analysis revealed substantial baseline variance among human quantifiers, particularly within the proton and carbon ion conditions for the BxPC-3 cell line. Under these conditions, surviving colonies exhibited significant morphological heterogeneity within single wells. This complexity naturally drives human disagreement, as manual scoring relies on subjective visual judgment that is easily fatigued by irregular structures. Besides, traditional manual scoring suffers from a lack of methodological standardization across different laboratories. Laboratories vary widely on whether counts are routinely verified via microscopy, evaluated by multiple blind observers, or estimated by a single individual. This methodological variability makes inter-laboratory comparisons fundamentally unreliable. The automated AI pipeline overcomes this bottleneck by applying consistent, pixel-level evaluation criteria. By removing the elements of cognitive fatigue, subjective bias, and variable laboratory protocols, the model ensures highly reproducible quantification and standardizes analysis even in the presence of the aberrant morphologies where human experts naturally diverge.

Besides mitigating the subjectivity of manual counting, reducing the processing time of the clonogenic assay was another major focus of this optimization. Traditional manual quantification is labor-intensive and completing a full experimental dataset manually could extend over several weeks. To achieve an automated pipeline capable of rapid, high-throughput quantification, we evaluated several acquisition platforms which were scored based on their acquisition speed and ability to capture whole well images. The EVOS M7000 system, equipped with an automated stage and 2x objective, proved significantly faster than alternative platforms. The EVOS M7000 reduced the acquisition time for a complete 6-well plate to three minutes, representing roughly 50% - 60% reduction in scanning time compared to competing systems. The AI-integrated segmentation pipeline further streamlined the workflow. It should be noted that while batch processing can take several days depending on the volume of samples, processing steps included, and local computer specifications, the system operates continuously without requiring human intervention. This operational efficiency is particularly advantageous in radiobiological research, where the inclusion of varying radiation qualities, doses, cell lines, and necessary statistical replicates rapidly inflates sample sizes.

To maintain this computational efficiency, we configured our segmentation using a semantic rather than an instance-based algorithm. While both approaches are available within the Zeiss Arivis Cloud platform, their practical utility was remarkable different. The semantic algorithm achieved robust results within few training iterations (3 for BxPC-3 and 7 for PANC-1). In contrast, the instance algorithm consistently underperformed, generating substantial false negatives. Unlike pixel-based semantic segmentation, instance segmentation attempts to identify and separate objects based on unique defining characteristics such as shape, size, and color ^11 12^. Because mammalian cell colonies exhibit vast morphological diversity, even within a single cell line, this likely reduced the instance algorithm’s generalizability, driving up both the false negative rate and necessary training time. However, because semantic segmentation classifies pixels rather than isolating distinct objects, an additional image processing step to separate merged colonies had to be implemented downstream in the pipeline.

While the transition to a semantic segmentation algorithm offered significant computational advantages, optimizing the imaging and training workflows was critical to achieving robust performance. First, optimizing initial plating densities to minimize colony merging reduces the reliability on post-processing to separate overlapping structures. Second, during acquisition, precise lighting calibration was necessary to ensure uniform background illumination and increase contrast between colonies and background. Additionally, to establish ground truth for training the models, we utilized a tablet-based system, which supported accurate, pixel-level manual contouring of colony edges. During the training phase, certain colony morphologies were difficult for the AI to delineate. For example, difference in cellular density between colonies resulted in false negatives for very sparsely populated colonies and colonies touching the well border. To account for this, during the next training iteration, selective annotations of those types of colonies were included. Training was deemed satisfactory when visual inspection showed satisfactory contouring of all colonies.

Besides the necessity of additional downstream processing, a broader limitation of the presented framework is the reliability on specialized infrastructure. Acquiring and processing whole-well images generates large datasets that demand digital storing and specialized computational power. Furthermore, the current pipeline relies on proprietary, commercial software, which carries licensing costs. Together, these limitations restrict accessibility for lower-resource laboratories. To overcome these limitations, future efforts should focus on enabling the integration of these trained deep learning segmentation algorithms into widely available, open-source image analysis platforms.

In addition, generalizability of the AI model to other cell lines remains a critical hurdle. While the semantic models developed for the specific morphologies of the PANC-1 and BxPC-3 cell lines demonstrated high accuracy, these deep learning architectures are inherently biased towards their training data. Applying these specific models to cell lines with different growth patterns, or images acquired with different systems will likely yield suboptimal segmentation outcomes. Our future work is addressing this generalizability issue by developing a singular, universal segmentation model trained on a diverse dataset of distinct cell lines. A major advantage of using cloud-based platforms like Zeiss Arivis Cloud is the possibility of direct shareability data. Researchers across different institutions can easily add and share images of their clonogenic experiments, thereby extending the diversity of the training dataset, and exchange and validate segmentation models, fostering a more collaborative, open-science approach for standardizing the clonogenic assay.

In conclusion, this study validates a deep learning-based semantic segmentation pipeline that significantly enhances the throughput, objectivity, and analytical depth of the clonogenic assay. By operating within the bounds of human consensus, the framework eliminates subjective visual bias and standardizes the derivation of radiobiological parameters, including relative biological effectiveness. Most importantly, transcending binary colony counts to include continuous morphological profiling provides a powerful new metric for evaluating complex cellular adaptation to different radiation qualities.

## Data Availability

The dataset and analysis pipelines in this study are available from the corresponding author on reasonable request.

## Acknowledgements

Support for this work was provided by the State of Florida Casey DeSantis funds to Mayo Clinic in Florida

## Author Contributions

W.E.R., J.M.D., and C.J.B. contributed to the conception of the experimental outline and methodology. W.E.R., A.K., K.R., and M.B. contributed to experimental procedures and data acquisition. W.E.R. performed data analysis and drafted the manuscript. All authors reviewed the manuscript.

## Literature

1. Franken, N. A., Rodermond, H. M., Stap, J., Haveman, J. & van Bree, C. Clonogenic assay of cells in vitro. Nat Protoc 1, 2315–2319 (2006). 10.1038/nprot.2006.339

2. Joiner, M. C. in Basic Clinical Radiobiology (ed M. C.; van der Kogerl Joiner, A.J.) Ch. 4, 32–42 (CRC Press, 2025).

3. Stone, H. B. et al. Preclinical Data on Efficacy of 10 Drug-Radiation Combinations: Evaluations, Concerns, and Recommendations. Transl Oncol 9, 46–56 (2016). 10.1016/j.tranon.2016.01.002

4. Brix, N., Samaga, D., Belka, C., Zitzelsberger, H. & Lauber, K. Analysis of clonogenic growth in vitro. Nat Protoc 16, 4963–4991 (2021). 10.1038/s41596-021-00615-0

5. Bray, M. A., Vokes, M. S. & Carpenter, A. E. Using CellProfiler for Automatic Identification and Measurement of Biological Objects in Images. Curr Protoc Mol Biol 109, 14.17.11–14.17.13 (2015). 10.1002/0471142727.mb1417s109

6. Choudhry, P. High-Throughput Method for Automated Colony and Cell Counting by Digital Image Analysis Based on Edge Detection. PLOS ONE 11, e0148469 (2016). 10.1371/journal.pone.0148469

7. Gomes, N. P., Frederick, B., Jacobsen, J. R., Chapnick, D. & Su, T. T. A High Throughput Screen with a Clonogenic Endpoint to Identify Radiation Modulators of Cancer. Radiat Res 199, 132–147 (2023). 10.1667/rade-22-00086.1

8. Zhang, L. Machine learning for enumeration of cell colony forming units. Vis Comput Ind Biomed Art 5, 26 (2022). 10.1186/s42492-022-00122-3

9. Makrai, L. et al. Annotated dataset for deep-learning-based bacterial colony detection. Sci Data 10, 497 (2023). 10.1038/s41597-023-02404-8

10. Denbeigh, J. M. et al. Clonogenic survival and relative biological effectiveness of pancreatic ductal adenocarcinoma under pristine and spread-out Bragg peak irradiation with protons and carbon ions. bioRxiv, 2026.2009.2030.754978 (2026). 10.64898/2026.09.30.754978

11. Cheng, B., Misra, I., Schwing, A. G., Kirillov, A. & Girdhar, R. in Proceedings of the IEEE/CVF conference on computer vision and pattern recognition. 1290–1299. 12 Zeiss. (2023).

